# Plastid Osmotic Stress influences cell differentiation at the Plant Shoot Apex

**DOI:** 10.1101/039115

**Authors:** Margaret E. Wilson, Matthew Mixdorf, R. Howard Berg, Elizabeth S. Haswell

## Abstract

The balance between proliferation and differentiation in the plant shoot apical meristem is controlled by regulatory loops involving the phytohormone cytokinin and stem cell identity genes. Concurrently, cellular differentiation in the developing shoot is coordinated with the environmental and developmental status of plastids within those cells. Here we employ an *Arabidopsis thaliana* mutant exhibiting constitutive plastid osmotic stress to investigate the molecular and genetic pathways connecting plastid osmotic stress with cell differentiation at the shoot apex. *msl2 msl3* mutants exhibit dramatically enlarged and deformed plastids in the shoot apical meristem, and develop a mass of callus tissue at the shoot apex. Callus production in this mutant requires the cytokinin receptor AHK2 and is characterized by increased cytokinin levels, down-regulation of cytokinin signaling inhibitors ARR7 and ARR15, and induction of the stem cell identity gene *WUSCHEL*. Furthermore, plastid stress-induced apical callus production requires elevated plastidic ROS, ABA biosynthesis, the retrograde signaling protein GUN1, and ABI4. These results are consistent with a model wherein the cytokinin/WUS pathway and retrograde signaling control cell differentiation at the shoot apex.

**SUMMARY STATEMENT:** Plastid osmotic stress influences differentiation at the plant shoot apex. Two established mechanisms that control proliferation, the cytokinin/WUSCHEL stem cell identity loop and a plastid-to-nucleus signaling pathway, are implicated.

## INTRODUCTION

The development of land plants provides a unique opportunity to study how cell differentiation is determined, as plant cell identity is highly plastic (Gaillochet and Lohmann, 2015). A classic illustration of plant cell pluripotency is the ability to produce a mass of undifferentiated cells referred to as callus. In nature, callus production is triggered by wounding or exposure to pathogens (Ikeuchi et al., 2013). In the laboratory, callus is typically induced by exogenous treatment with two phytohormones, cytokinin (CK) and auxin (Skoog and Miller, 1957).

Callus is frequently derived from meristem or meristem-like cells (Jiang et al., 2015; Sugimoto et al., 2011). Meristems are small self-renewing pools of undifferentiated cells from which new organs are derived as the plant grows (Aichinger et al., 2012; Gaillochet and Lohmann, 2015). Above ground, the maintenance and regulation of the shoot apical meristem (SAM) is critical for the proper specification and positioning of leaves (Barton, 2010). SAM identity requires both the imposition of stem cell fate by the WUSCHEL(WUS)/CLAVATA(CLV) signaling circuit (Fletcher et al., 1999; Schoof et al., 2000) and the suppression of differentiation by SHOOTMERISTEMLESS (STM) (Endrizzi et al., 1996; Long et al., 1996).

CK plays a key role in the function of the SAM. In the central zone (CZ), CK promotes proliferation, while auxin promotes differentiation in the peripheral zone (PZ) (Schaller et al., 2015). Localized CK perception and response specifies the organizing center of the SAM, also the region of *WUS* expression (Chickarmane et al., 2012; Gordon et al., 2009; Zurcher et al., 2013). WUS activity maintains itself through a positive feedback loop involving CK response via Type-A Arabidopsis Response Regulators (ARRs), key negative regulators of CK signaling (Leibfried et al., 2005; Schuster et al., 2014; To et al., 2007; Zhao et al., 2010). In the SAM, auxin acts to repress *ARR7* and *ARR15*, (Zhao et al., 2010). Thus, auxin and CK synergize to regulate the core WUS/CLV pathway, maintaining a balance between differentiation and proliferation in the SAM (Gaillochet and Lohmann, 2015; Ikeuchi et al., 2013).

Shoot development also depends on plastids, endosymbiotic organelles responsible for photosynthesis, amino acid, starch, and fatty acid biosynthesis and the production of many hormones and secondary metabolites (Neuhaus and Emes, 2000). As cells leave the SAM and take on the appropriate cell identity within leaf primordia, the small, undifferentiated plastids, called proplastids, inside them must also differentiate, usually into chloroplasts. Many mutants lacking functional plastid-localized proteins exhibit secondary defects in leaf cell specification ((Larkin, 2014; Luesse et al., 2015; Moschopoulos et al., 2012), providing genetic evidence that normal leaf development depends upon plastid homeostasis. The integration of plastid differentiation into the process of development likely requires tightly regulated and finely tuned two-way communication between the plastid and the nucleus, including both anterograde (nucleus-to-plastid) and retrograde (plastid-to-nucleus) signaling. An increasing number of overlapping retrograde signaling pathways, triggered by developmental or environmental defects in plastid function, have been identified (Chan et al., 2015; Woodson and Chory, 2012). Numerous retrograde signals have been proposed, including intermediates in isoprenoid (Xiao et al., 2012) biosynthesis, heme (Woodson et al., 2011), phosphonucleotides (Estavillo et al., 2011), reactive oxygen species (ROS) (Wagner et al., 2004) and oxidized carotenoids (Ramel et al., 2012). Despite the diversity, all these pathways and signals link a disruption in plastid homeostasis to altered nuclear gene expression (Chan et al., 2015).

One retrograde signaling pathway that may serve to connect plastid signals to shoot development is the GENOMES UNCOUPLED (GUN1) / ABA-INSENSITIVE 4 (ABI4) pathway (Fernandez and Strand, 2008; Leon et al., 2012). ABI4 is a nuclear transcription factor involved in many plant developmental pathways, including response to ABA, sugar signaling, and mitochondrial retrograde signaling (Leon et al., 2012). GUN1 is a plastid protein of unclear molecular function thought to function with ABI4 in at least two retrograde signaling pathways, one that coordinates plastid and nuclear gene expression during development, and one that respond to defects in chlorophyll biosynthesis (Cottage et al., 2010; Koussevitzky et al., 2007; Sun et al., 2011).

We have been using an Arabidopsis mutant with constitutively osmotically stressed plastids (*msl2 msl3* (Veley et al., 2012)) as a model system to address the developmental effects of plastid dysfunction. MSL2 and MSL3 are two members of the MscS-Like family of mechanosensitive ion channels that localize to the plastid envelope and are required for normal plastid size, shape, and division site selection (Haswell and Meyerowitz, 2006; Wilson et al., 2011). By analogy to family members in bacteria (Levina et al., 1999) and plants (Hamilton et al., 2015), and based on *in vivo* experiments (Veley et al., 2012), MSL2 and MSL3 are likely to serve as osmotic “safety valves”, allowing plastids to continuously maintain osmotic homeostasis during normal growth and development. In addition, *msl2 msl3* mutant plants exhibit small stature, variegated leaf color, and ruffled leaf margins. These whole-plant defects can be attributed to plastid osmotic stress, as they are suppressed by environmental and genetic manipulations that increase cytoplasmic osmolarity and draw water out of the plastid (Veley et al., 2012; Wilson et al., 2014). Here we report a new and unexpected phenotype associated with *msl2 msl3* mutants, and establish the molecular and genetic pathways that underlie it.

## RESULTS

### *msl2 msl3* double mutants develop callus at the shoot apex

When grown on solid media for over 14 days, *msl2 msl3* double mutant plants developed a proliferating mass of undifferentiated cells, or callus, at the apex of the plant (Fig. 1A-D). A single mass at the center of the apex, twin masses associated with the cotyledon petioles, or occasionally three masses were observed Fig. 1A-C). New green tissue was frequently observed growing out of the apical callus (white arrow, Fig. 1D). This phenotype was not observed in soil-grown plants.

**Fig. 1.**
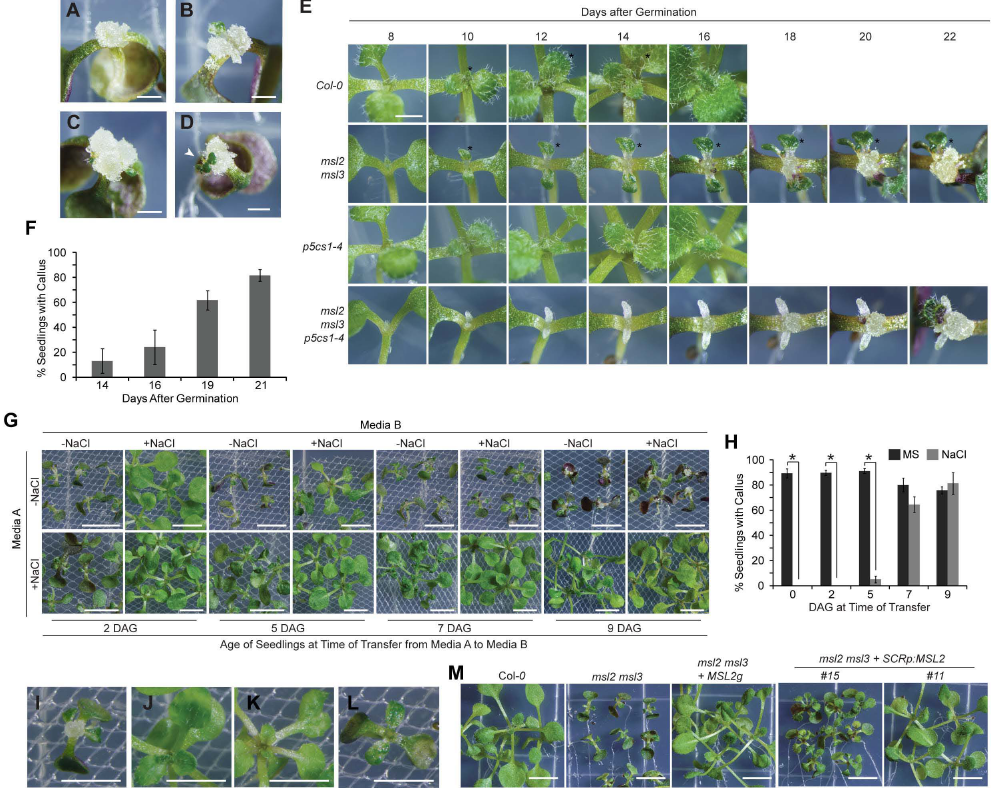
***msl2 msl3* double mutants develop callus at the shoot apex.** (A-D) Bright-field images of 21-day-old *msl2 msl3* seedlings grown on solid media. Shooty callus is indicated with a white arrowhead in (D). (E) Time course of callus formation. (F) Average percent of *msl2 msl3* seedlings with visible callus at the indicated time after germination. Five plates of seedlings, n > 10 seedlings per plate, were analyzed per time point. (G) 21-day-old seedlings scored in (H) (top) or seedlings transferred from MS with 82 mM NaCl to MS at indicated time points (bottom). (H) The average percent of *msl2 msl3* seedlings showing visible callus at 21 DAG after transfer from MS to MS (black bars) or from MS to MS with 82 mM NaCl (grey bars) at the indicated time points. 3 plates of seedlings, n > 10 seedlings per plate, were analyzed per treatment. Error bars indicate SEM. *, difference between indicated samples, *P* < 0.01 (Student’s *t*-test). (I-M) Close-up images of individual seedlings shown in (G) grown without NaCl (I) or transferred to NaCl plates at 2 (J), 5 (K), or 7 (L) DAG. (M) Bright-field images of 14-day-old *msl2 msl3* seedlings harboring the *MSL2* transgene (*MSL2g*) or *MSL2* under the control of the *SCR* promoter. Size bars (A-E), 1 mm, (G, M), 5 mm, (I-L), 2.5 mm.

Masses of callus tissue were apparent to the naked eye at the apex of *msl2 msl3* seedlings between 14 and 16 days after germination (DAG) and continued to grow in size through 22 DAG (Fig. 1E, top two rows). As previously documented (Haswell and Meyerowitz, 2006; Jensen and Haswell, 2012; Wilson et al., 2014), *msl2 msl3* leaves were small and malformed. To facilitate comparison, the same leaf from each genotype is marked with an asterisk in the time course in Fig. 1E. The percent of *msl2 msl3* seedlings with callus increased to ~82% by 21 DAG (Fig. 1F); callus was not observed in any wild type plants at any developmental stage.

We previously showed that growth on media containing increased levels of osmolytes (sugars or salt) suppressed the plastid morphology and leaf development phenotypes of *msl2 msl3* mutants, likely by increasing cytoplasmic osmolarity and reducing plastid hypoosmotic stress (Veley et al., 2012; Wilson et al., 2014). To determine if the same were true for callus formation, and to assess the dependence of suppression on age and developmental stage, seedlings were germinated on MS solid media and transferred to media containing 82 mM NaCl at 2, 5, 7, or 9 DAG and assessed at 21 DAG for callus formation and leaf development. Full suppression of callus formation and abnormal leaf development was only observed in seedlings transferred to solid media containing NaCl from MS at 2 DAG (Fig. 1G-J), establishing a small developmental window after which alleviation of plastid hypoosmotic stress can no longer suppress leaf morphology defects and apical callus production in *msl2 msl3* mutants. Suppression was also observed when seedlings were grown on media containing sucrose, sorbitol or mannitol, indicating that the effect is osmotic (Fig S1A.) The converse experiment, in which seedlings were transferred from salt-containing media to normal MS showed that plastid osmotic homoeostasis is continuously required to maintain normal leaf development (Fig. 1H and S1B).

### MSL2 function is required in the SAM or leaf primordia to prevent callus production

To determine if MSL2 expression in the SAM and leaf primordia is required to prevent callus production in the *msl2 msl3* background, we examined apical callus production in *msl2 msl3* mutant plants expressing *MSL2* from the *SCARECROW (SCR)* promoter, which is expressed in the L1 cell layer of the meristem and leaf primordia (Wysocka-Diller et al., 2000). As shown in Fig. 1H, apical callus formation was fully suppressed in T2 *msl2 msl3 SCRpMSL2* lines. While these plants developed multiple sets of developmentally normal true leaves, the majority of T2 lines examined exhibited reduced stature.

### Cells and plastids at the apex of young *msl2 msl3* mutants are morphologically abnormal

Closer examination of the apex of young (4-day-old) *msl2 msl3* mutant seedlings revealed a cluster of disorganized and heterologously shaped cells with low electron density at the expected location of the SAM (Fig 2A). Most cells within the cluster contained large, spherical, clear entities. In comparison, the SAM cells of a wild type seedling were organized into cell layers and were electron-dense. We next used transmission electron microscopy to further characterize the morphology of developing chloroplasts and proplastids in the SAM and surrounding tissue of *msl2 msl3* mutant seedlings (Fig 2B). In the *msl2 msl3* mutant, young chloroplasts were enlarged, lacked the lens shape of wild-type chloroplasts, and exhibited a disorganized developing thylakoid network (Fig. 2B-D and S2B). In agreement with previous reports (Charuvi et al., 2012), wild-type proplastids appeared as small structures (0.5-1 μm in diameter) containing rudimentary thylakoid networks of varying developmental stages, plastoglobules, and double membranes (Fig. 2G and Fig. S2A). However, in the SAM of *msl2 msl3* mutants, entities with these established features had decreased stromal density and were greatly enlarged compared to the wild type (asterisks in Fig. 2A and S2B, Fig. 2E). Many exceeded 5 μm in diameter. All were clearly bound by a double membrane (white arrow, Fig. 2F and S2D-E). These data show that the greatly enlarged phenotype of non-green plastids of the leaf epidermis (Haswell and Meyerowitz, 2006; Veley et al., 2012) extends to proplastids of the SAM.

**Fig. 2.**
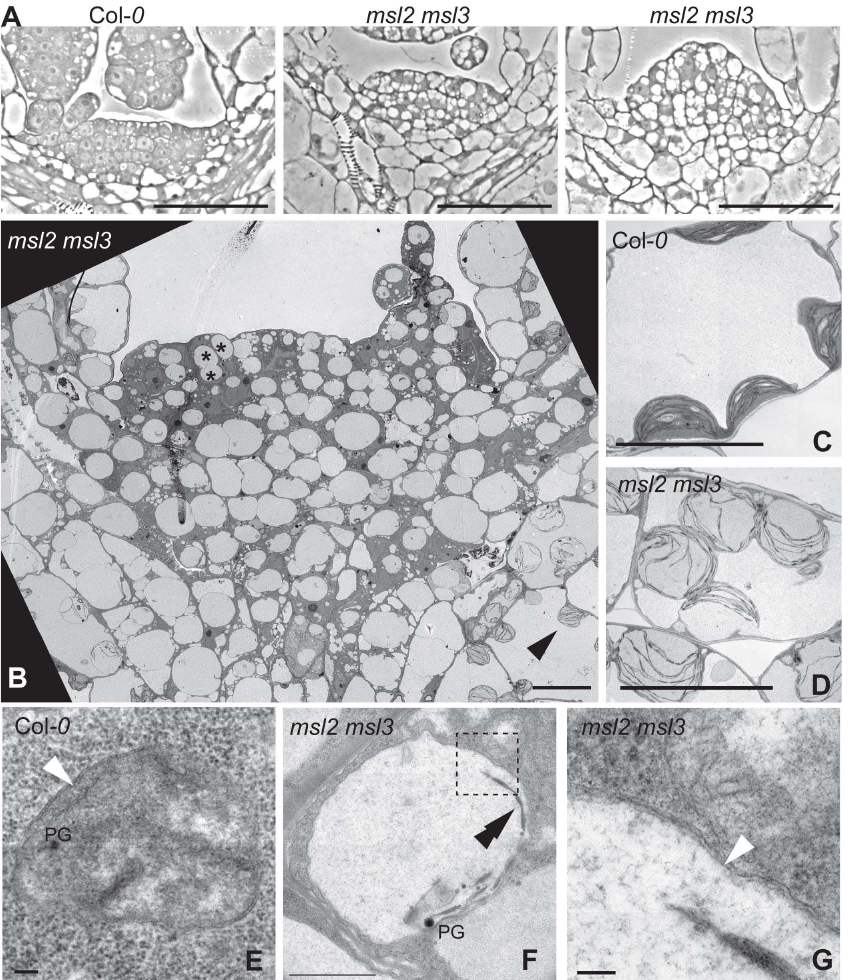
**Four-day-old *msl2 msl3* mutants exhibit abnormal cellular organization and plastids at the SAM.** (A) Phase contrast images of the SAM region of seedlings of the indicated genotypes. Size bar, 50 μm. (B-G) Transmission electron microscope (TEM) images of *msl2 msl3* mutant SAM (B) and developing chloroplasts (C-D) and proplastids (E-F) in wild type and *msl2 msl3* mutant seedlings. (G) Magnified region of *msl2 msl3* plastid envelope, indicated by box in F. Examples of proplastids and developing chloroplasts are indicated in *msl2 msl3* mutant by asterisks and black arrowheads, respectively. Size bar, 10 μm (B-D), 1 μm (E), and 100 nm (F-G). Plastoglobules (PG), double membrane (white arrowheads), and thylakoids (double black arrowheads) are indicated.

### Callus produced in *msl2 msl3* mutants is associated with an altered cytokinin: auxin ratio and requires CK perception

The production of shooty callus has been associated with increased production or availability of CKs and an imbalance in the cytokinin: auxin ratio (Frank et al., 2002; Frank et al., 2000; Lee et al., 2004). Consistent with these observations, apical callus production, dwarfing and leaf phenotypes of the *msl2 msl3* double mutant were strongly suppressed when seedlings were grown on media with the synthetic auxin 1-naphthaleneacetic acid (NAA) (Fig. 3A, C). The leaf epidermis of *msl2 msl3* seedlings contain grossly enlarged and round non-green plastids, as visualized with the fluorescent plastid marker RecA-dsRED (Haswell and Meyerowitz, 2006; Veley et al., 2012; Wilson et al., 2014). This phenotype was not altered by growth on 2 *μ*M NAA (Fig. 3B), indicating that NAA treatment suppresses callus formation downstream of effects on plastid morphology. In agreement with previous data, growth on media supplemented with osmolytes fully suppressed the plastid morphology defects (Veley et al., 2012). Trans-Zeatin-riboside levels were increased ~6.5-fold in *msl2 msl3* mutant seedlings compared to the wild type, but IAA levels were not significantly changed (Fig. 3D).

**Fig. 3.**
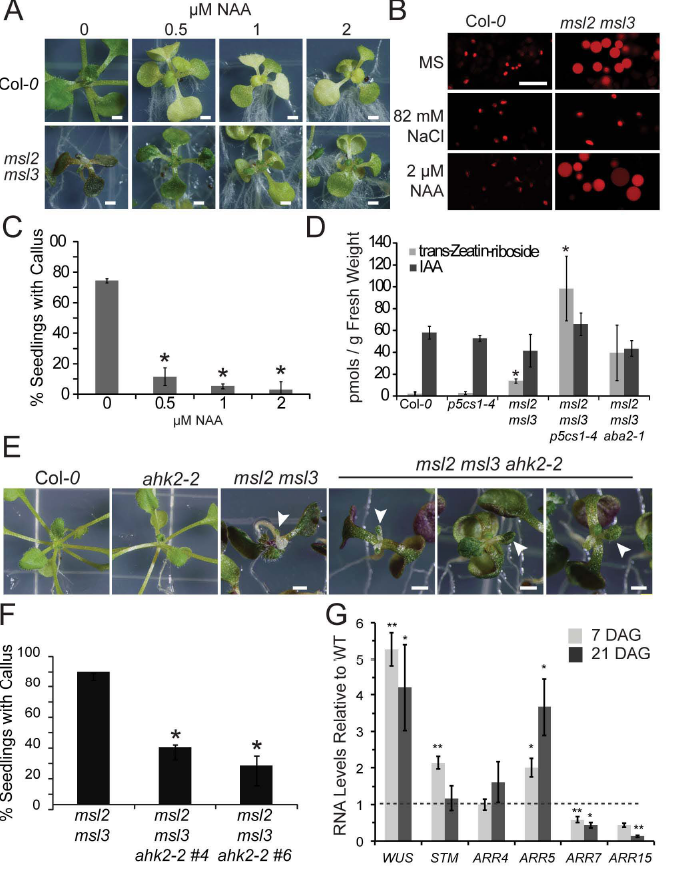
**Callus produced in *msl2 msl3* mutants is associated with increased CK production and requires CK signaling.** (A) Seedlings grown for 21 days on solid media containing the indicated concentration of NAA. Size bar, 1 mm. (B) Confocal micrographs of non-green plastids in the first true leaf of *msl2 msl3* mutants harboring the RecA-dsRED plastid marker and grown on indicated media. Size bar, 10 μm. (C) Percent of *msl2 msl3* mutants exhibiting callus when grown on NAA. The average of three biological replicates of ≥ 20 seedlings each is presented. Statistical groups were determined by ANOVA followed by Tukey’s HSD test, P < 0.01. (D) Trans-Zeatin-riboside and IAA levels of 21-day-old seedlings determined by liquid chromatography-mass spectrometry/mass spectrometry analysis as in (Chen et al., 2009). The average of three biological replicates of ≥ 30 seedlings each is presented. Error bars indicate SD, *, difference from wild type, *P* < 0.01 (Student’s *t*-test). Images of seedlings (E) and callus production rates (F) of *msl2 msl3 ahk2-2* triple mutant and parental lines. Size bar, 1 mm. White arrows mark deformed leaves. The average of four biological replicates of ≥ 20 seedlings each is presented. Error bars indicate SEM. *, difference from *msl2 msl3*, *P* < 0.01 (Student’s *t*-test). (G) Quantitative RT-PCR analysis of gene expression in the *msl2 msl3* mutant. The average of three biological replicates (two technical replicates; n ≥ 25 seedlings each) is presented. Error bars indicate SEM, asterisks indicate difference from wild type of same age, **P ≤ 0.05, **P ≤ 0.01 (Student’s *t*-test).

The *ARABIDOPSIS HISTIDINE KINASE (AHK)2* gene encodes one of three related histidine kinases known to function as CK receptors in Arabidopsis (Ueguchi et al., 2001; Yamada et al., 2001), and an *AHK2* loss of function mutant, *ahk2-2* (Higuchi et al., 2004) is impaired in CK-induced up-regulation of *WUS* (Gordon et al., 2009). Two independently isolated *msl2 msl3 ahk2-2* triple mutants showed a significant reduction in apical callus formation with less than 40% of seedlings developing callus (Fig. 3F). However, the *msl2 msl3 ahk2-2* triple mutants were similar to the *msl2 msl3* double with respect to leaf developmental defect (Fig. 3E). Incomplete suppression may be due to redundancy among CK receptors (Gordon et al., 2009); or callus formation and leaf morphology defects may be produced through different pathways.

To determine if feedback loops involving *WUS*, *ARRs* (Gordon et al., 2009; Schuster et al., 2014), or the meristem identity gene *STM* (Scofield et al., 2013) were mis-regulated in *msl2 msl3* seedlings, quantitative RT-PCR was used to determine transcript levels in the aerial tissue of 7- and 21-day-old seedlings (Fig. 3G). In *msl2 msl3* mutant seedlings *WUS* transcript levels were up-regulated 5.3-fold compared to the wild type by 7 DAG (prior to visible callus development, Fig. 1E); and 4-fold by 21 DAG. *STM* expression levels were up-regulated 2.1-fold in 7-day-old *msl2 msl3* mutants, but were not distinguishable from the wild type at 21 DAG. Consistent with previous observations that they inhibit callus formation (Buechel et al., 2010; Liu et al., 2016), *ARR7* and *ARR15* transcript levels were reduced in both 7 and 21 day old *msl2 msl3* mutant seedlings compared to wild type, exhibiting 2 to 2.6-fold and 3- to 6-fold decreases, respectively. Transcriptional repression is specific to *ARR7* and *ARR15* as transcript levels of two other A-type *ARR* genes, *ARR4* and *ARR5,* were elevated in *msl2 msl3* mutant seedlings compared to wild type. Hypocotyls from *msl2 msl3* mutants did not efficiently produce callus in vitro, indicating that an additional signal or signals are required to produce callus outside the SAM in these mutants (Fig. S3A). Furthermore, multiple genes involved in wound-inducible and auxin-inducible callus production show altered expression in *msl2 msl3* mutants, including *WIND1*, *WIND3*, *KPR2*, *KPR7*, *TSD1* and *TSD2* (Anzola et al., 2010; Frank et al., 2002; Iwase et al., 2011; Krupkova and Schmulling, 2009) (Fig. S3B, C).

### Preventing Pro biosynthesis results in a dramatic increase in callus formation and CK levels in the *msl2 msl3* background

It was not obvious how the constitutive plastid osmotic stress experienced by *msl2 msl3* mutants might elicit these effects, but we reasoned that the hyper-accumulation of solutes previously observed in *msl2 msl3* mutants, especially the compatible osmolyte Proline (Pro) (Wilson et al., 2014), might be responsible. To test this hypothesis, we crossed the *pyrroline-5-carboxylate synthetase1-1* (*p5cs1-4*) lesion into the *msl2 msl3* mutant background. P5CS1 catalyzes the primary step in the inducible production of Pro (Verslues and Sharma, 2010), and stress-induced Pro levels are low in this mutant (Szekely et al., 2008). *msl2 msl3 p5cs1-4* triple mutant seedlings exhibited larger calluses than the *msl2 msl3* double, frequently forming multiple calluses (Figs. 1E, and S4A), and formation was more observed at earlier stages of development in *msl2 msl3 p5cs1-4* triple compared to *msl2 msl3* double mutants (Fig. 4A, light blue bars)—at 14 DAG over 40% of triple mutant seedlings had visible callus, while only 15% of the double *msl2 msl3* mutants did (Fig. 4A, light blue bars). Neither wild type nor *p5cs1-4* single mutants produced callus at any time point. A different mutant allele of *P5CS1*, *p5cs1-1* (Szekely et al., 2008), also enhanced callus formation in the *msl2 msl3* background (Fig. S4B-C). In addition *msl2 msl3 p5cs1-4* triple mutants exhibited 7 times more trans-Zeatin-riboside levels than *msl2 msl3* mutants (Fig. 3D). Supplementing growth media had no effect on callus production in *msl2 msl3* double or *msl2 msl3 p5cs1-4* triple mutants at any time point (Fig. 4A), even though *msl2 msl3 p5cs1-4* triple mutant seedlings grown on 20 mM Pro for 21 DAG contained as much Pro as the *msl2 msl3* double (Fig. 4B).

**Fig. 4.**
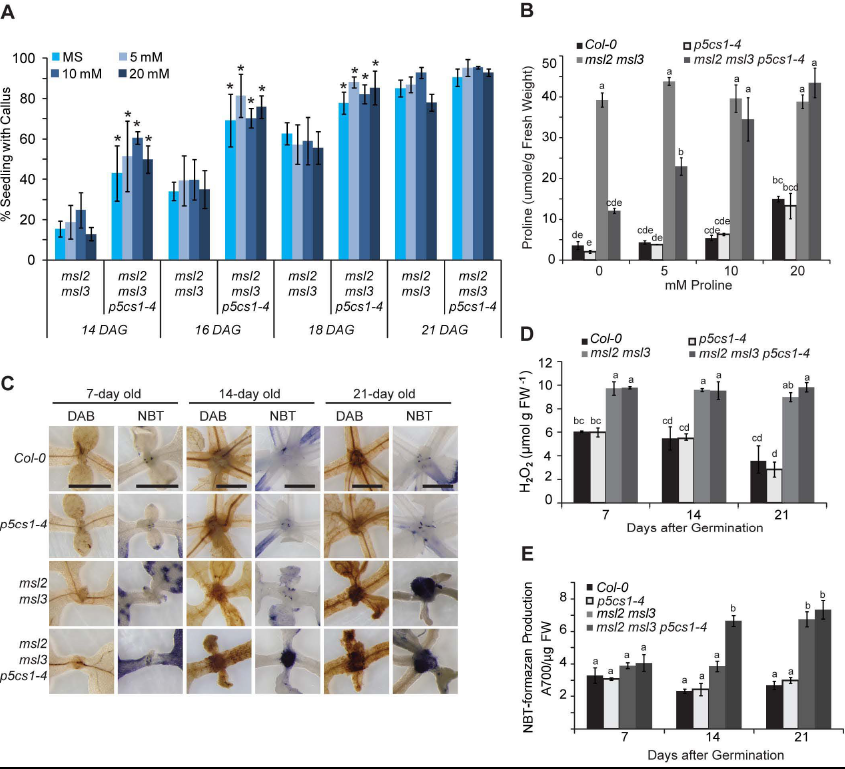
**Preventing Pro biosynthesis results in a dramatic increase in callus formation in the *msl2 msl3* background. (**A) Production of apical callus in *msl2 msl3* and *msl2 msl3 p5cs1-4* seedlings in the presence and absence of exogenously provided Pro. The average of three biological replicates is presented, *n* ≥ 25 seedlings each. Error bars represent SD. *, difference from *msl2 msl3* seedlings of the same age, *P* < 0.01 (Student’s *t-*test). (B) Pro content of the aerial tissue of mutant and wild-type seedlings grown as in (A), measured as in (Wilson et al., 2014). The average of two biological replicates (each performed in triplicate; n ≥ 30 seedlings each) is presented. (C) Images of the meristematic region of seedlings stained with DAB or NBT. Size bar, 1 mm. Quantification of H_2_O_2_ content by Amplex Red (D) and O_2_^-^ production by NBT-Formazan production (E) in mutant and wild-type seedlings of indicated age. The average of three biological replicates (*n* ≥ 20 seedlings each), two technical replicates each, is presented in both (D) and (E). Statistical grouping in (B), (D), and (E) was performed as in Fig. 3C, and error bars indicate SEM. FW, fresh weight.

Thus, it is not the absence of Pro itself that leads to the enhanced callus production in the *msl2 msl3 p5cs1-4* triple mutants. Instead, disrupting the process of Pro biosynthesis could enhance callus formation in the *msl2 msl3* mutant background if ROS accumulation were involved, as Pro biosynthesis is a key reductive pathway that helps maintain cellular redox homeostasis (Szabados and Savoure, 2010). To characterize the levels and localization of ROS in *msl2 msl3* double and *msl2 msl3 p5cs1-4* triple mutants, 7-, 14-, and 21-day-old mutant and wild-type seedlings were stained with 3, 3 diaminobenzidine (DAB) to detect hydrogen peroxide (H_2_O_2_) or nitroblue tetrazolium (NBT) to detect superoxide (O_2_^-^) (Fig. 4C). Both double and triple mutants accumulated high levels of H_2_O_2_ and O_2_^-^ in the SAM region relative to the *p5cs1-4* single mutant or the wild type. Precipitate was visible earlier in the *msl2 msl3 p5cs1-4* triple mutant (by 14 DAG compared to 21 DAG in the in the double *msl2 msl3* mutant). Quantitation of H_2_O_2_ levels using an Amplex Red enzyme assay at these same time points showed consistent accumulation of H_2_O_2_ in the double and triple mutants, rising to nearly three-fold over wild type by 21 DAG (Fig. 4D). Levels of formazan, the product of NBT reduction by O_2_^-^, increased in *msl2 msl3* and *msl2 msl3 p5cs1-4* to more than 2.5-fold above wild-type levels at 21 DAG (Fig. 4E). Calluses generated by incubating *Arabidopsis thaliana* Col*-0* root explants on Callus Inducing Media also showed strong NBT staining in areas of cell proliferation (Fig. S4), providing evidence that ROS accumulation may be a general feature of callus tissue (Lee et al., 2004).

### The accumulation of superoxide in response to plastid hypo-osmotic stress is required for callus formation in the *msl2 msl3* background

In plastids, the photoreduction of molecular O_2_ generates O_2_^-^, which is then rapidly converted into H_2_O_2_ by plastid-localized superoxide dismutase enzymes (Asada, 2006). To determine if plastid osmotic stress and the accumulation of O_2_^-^ at the apex of *msl2 msl3* double and *msl2 msl3 p5cs1-4* triple mutants play causative roles in callus formation, plants were germinated on or transferred to solid media with 82 mM NaCl or with TEMPOL (a O_2_^-^ scavenger) at 3 or 7 DAG and stained with NBT at 21 DAG. Growth on media containing NaCl suppressed O_2_^-^ accumulation in *msl2 msl3* double and *msl2 msl3 p5cs1-4* triple mutant seedlings (Fig. 5A), completely prevented callus formation (Fig. 5B), suppressed defects in non-green plastid morphology (Fig. 5C) and leaf morphology (Fig. 5A). Thus, plastid osmotic stress is responsible for O_2_^-^ accumulation as well as callus formation in *msl2 msl3* and *msl2 msl3 p5cs1-4* mutants. Consistent with Fig. 1H, growth on NaCl suppressed callus formation and ROS accumulation only if supplied prior to 7 DAG.

**Fig. 5.**
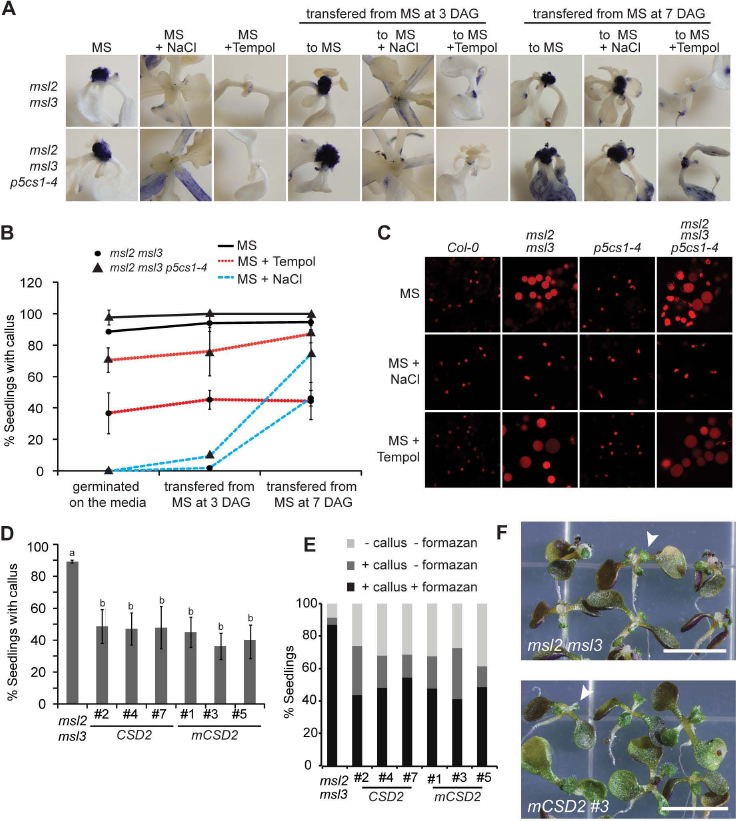
**Hyper-accumulation of superoxide is required for callus formation in *msl2 msl3* and *msl2 msl3 p5cs1* seedlings.** (A) NBT stained seedlings provided with TEMPOL or NaCl at 0, 3 or 7 DAG. Size bar, 1mm. (B) Callus production in mutant seedlings when grown on MS, MS with 1 mM TEMPOL or MS with 82 mM NaCl. (C) Confocal micrographs of non-green plastids in the first true leaf of *msl2 msl3* mutants harboring the RecA-dsRED plastid marker grown on the indicated media. Size bar, 10 μm. (D) Apical callus production in T2 lines segregating transgenes that over-express *CSD2* or *mCSD2*. The average of 4 biological replicates (*n* ≥ 35 seedlings each replicate) is presented. Error bars indicate SD. Statistical grouping was performed as in Fig. 3C. (E) Percent of seedlings from (D) stained with NBT accumulating formazan at the shoot apex. (*n* ≥ 20 seedlings). (F) Images of *CSD2* over-expression T2 lines. Size bar, 1 mm. White arrows mark deformed leaves.

Growth on TEMPOL-containing media successfully prevented the accumulation of O_2_^-^ in the SAM of *msl2 msl3* and *msl2 msl3 p5cs1-4* mutants, regardless of seedling age at time of application (Fig. 5A), but did not affect plastid morphology (Fig, 5C) nor leaf morphology defects (Fig. 5A). Treating double and triple mutant seedlings with TEMPOL at all developmental stages partially suppressed callus formation (Fig. 5C). Less than 40% of double mutant seedlings exhibited apical calluses when grown in the presence of TEMPOL, compared to > 90% when grown on MS without TEMPOL. TEMPOL-mediated callus suppression is thus independent or downstream of the developmental window in which addition of osmotic support must be provide for complete suppression of aerial phenotypes (Figs 1H and 5A-B).

A complementary genetic approach to suppressing O_2_^-^ accumulation in the plastid was taken by over-expressing wild type or miR398-resistant forms of the chloroplast-localized Cu/Zn Superoxide Dismutase CSD2 (Kliebenstein et al., 1998; Sunkar et al., 2006) in the *msl2 msl3* mutant background. The percentage of seedlings with visible apical callus at 21 DAG was decreased to 50% or less in six independent T2 lines segregating either wild-type (*CSD2*, lines #2, #4, and #7) or miR398 resistant (*mCSD2*, lines #1, #3, #5) *CSD2* over-expression constructs (Fig. 5D). Approximately 50% of T2 seedlings showed strong O_2_^-^ accumulation by NBT staining, which was localized to developing callus, compared to 90% of *msl2 msl3* seedlings (Fig. 5E). 15-30% of seedlings in the CDS2 over-expression lines exhibited both low levels of O_2_^-^ accumulation at their shoot apex and did not develop callus; less than 5% of *msl2 msl3* seedlings had these characteristics. Over-expression of *CSD2* did not suppress abnormal leaf development (white arrows, Fig. 5F).

### Callus production requires ABA biosynthesis, ABI4 and GUN1

As *msl2 msl3* double mutants exhibit increased levels of ABA and up-regulation of ABA biosynthesis genes (Wilson et al., 2014), we hypothesized that a pathway involving the hormone ABA and the *GUN1* and *ABI4* gene products might account for the pleiotropic defects in the *msl2 msl3* mutant (Koussevitzky et al., 2007; Zhang et al., 2013). To determine if ABA biosynthesis was required for callus production, we assessed the effect of introducing the *abscisic acid-deficient 2-1* allele (Schwartz et al., 1997) on callus formation in the *msl2 msl3* background. Indeed, callus formation and leaf development defects were strongly suppressed in a triple *msl2 msl3 aba2-1* mutant, with only ~20% of seedlings developing callus at the shoot apex by 21 DAG (Fig. 6A, B). Similar suppression of *msl2 msl3* aerial defects was observed in four other independently isolated *msl2 msl3 aba2-1* triple mutant lines. The *msl2 msl3 aba2-1* triple mutant accumulated trans-Zeatin-riboside to levels similar to the *msl2 msl3* double mutant (Fig. 3D), suggesting that ABA and CK promote callus formation through independent pathways.

**Fig. 6.**
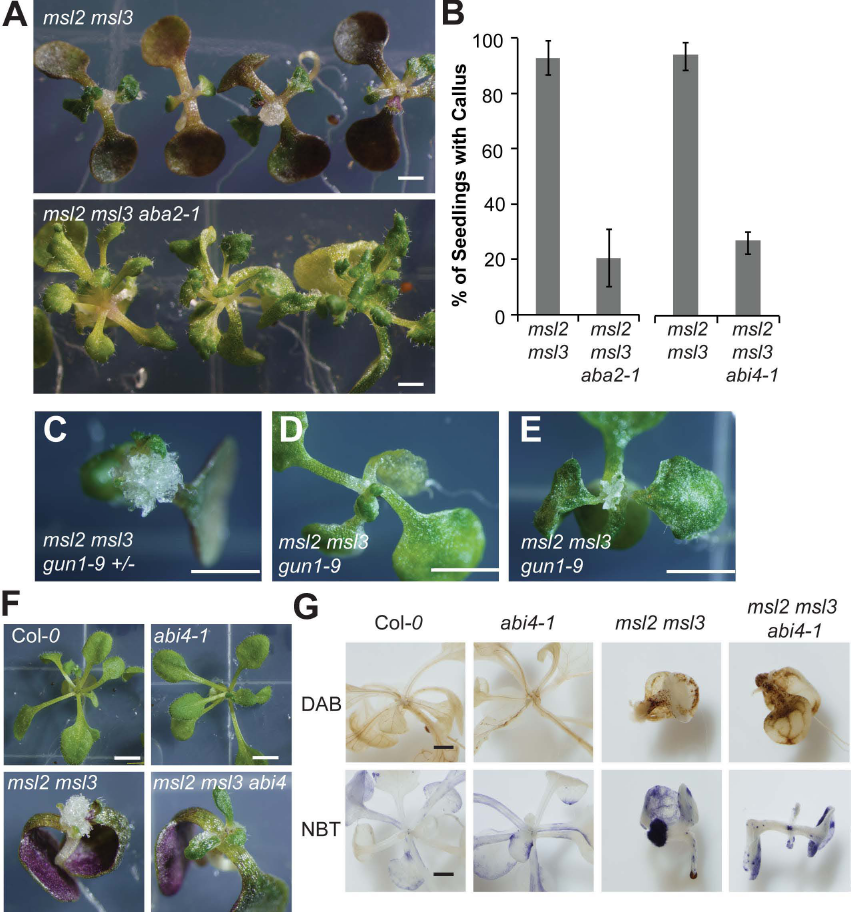
**Callus production requires ABA biosynthesis, GUN1 and ABI4.** (A) 21-day-old *msl2 msl3* double and *msl2 msl3 aba2-1* triple mutant seedlings. (B) Callus production in higher order mutants at 21 DAG. The average of 3 biological replicates (*n* ≥ 15 seedlings per replicate) is presented. Error bars indicate SEM. (C-D) 21-day-old *msl2 msl3 gun1-9* (+/−) (C) and *msl2 msl3 gun1-9* (−/−) (D, E) siblings at 21 DAG. (F) 21-day-old *msl2 msl3 abi4-1* seedlings with relevant parental controls. (G) 21-day-old seedlings of the indicated genotypes stained with DAB or NBT. (A, F, G) Size bar, 1 mm. (C, D, E) Size bar, 2.5 cm.

To test if GUN1 is involved in the perception of signals generated by plastid osmotic stress, we crossed the *gun1-9* (Koussevitzky et al., 2007) allele into the *msl2 msl3* mutant background and analyzed the offspring of a single *msl2 msl3 gun1-2 (+/−)* mutant plant. Of twenty-seven triple *msl2 msl3 gun1-9* seedlings identified by PCR genotyping, none formed apical callus when grown on solid media, whereas 16 of 19 genotyped sibling *msl2 msl3 gun1-9* (+/−) seedlings did. In addition, *msl2 msl3 gun1-9* triple mutant siblings produced larger, greener, and more normally shaped true leaves than their *msl2 msl3* and *msl2 msl3 gun1-9+/1* siblings (Fig. 6C, D). In some seedlings, small and chlorotic true leaves developed at the seedling apex (Fig. 6E, S5).

The strong *abi4-1* allele (Finkelstein et al., 1998) was also introduced into the *msl2 msl3* background, and *msl2 msl3 abi4-1* mutants also exhibited reduced apical callus formation; only ~26% of *msl2 msl3 abi4-1* seedlings from two independently isolated lines developed callus (Fig. 6F). Neither leaf developmental defects nor ROS accumulation was suppressed in the *msl2 msl3 abi4-1* mutant, and *msl2 msl3 abi4-1* seedlings stained with DAB or NBT showed a pattern of ROS accumulation similar to the *msl2 msl3* double mutant (Fig. 6F, G). These staining patterns were not observed in wild type or *abi4-1* single mutants. These results are consistent with a model wherein ABI4 functions downstream of ROS accumulation, and likely in a pathway with GUN1, to induce apical callus formation in response to plastid osmotic stress.

## DISCUSSION

One of the most fundamental decisions a cell can make is whether to proliferate or to differentiate. In the plant SAM, this decision must be spatially and temporally controlled, so that cells remain in an undifferentiated, pluripotent state in the central zone of the meristem and then differentiate properly as they are recruited into organs at the PZ. Here we use the *msl2 msl3* mutant as a model system to show that, in the Arabidopsis SAM, proliferation versus differentiation signals are coordinated not only at the tissue- and cellular-level, but also at the organellar level. We further applied genetic, molecular, biochemical, and pharmacological approaches to identify two non-redundant pathways through which plastid osmotic stress may produce apical callus (illustrated in Fig. S6).

### Plastid osmotic stress results in the production of callus at the plant SAM

Plants lacking functional versions of the mechanosensitive ion channel homologs MSL2 and MSL3 robustly produce callus tissue at the apex of the plant when grown on solid media (Fig. 1). Molecular complementation with *MSL2* under the control of the *SCARECROW* promoter established that MSL2/MSL3 are required only in the L1 layer of the CZ and/or PZ of the SAM to prevent callus formation (Fig. 2G) and that their function is critical during the first 5 days after germination (Fig 1G, H). The production of apical callus in the *msl2 msl3* mutant is associated with dramatically enlarged and developmentally abnormal plastids specifically in the SAM (Fig. 2A-F, S2A-D). Developmentally defective plastids were also observed in the SAM of another apical callus-producing mutant, *tumorous shoot development (tsd)1* (Frank et al., 2002).

### Increased proliferation and the production of callus in *msl2 msl3* mutants is associated with a disruption in the CK/WUS feedback loop

The *msl2 msl3* mutants exhibit several previously established hallmarks of increased proliferation at the SAM, including increased levels of CK, up-regulation of the stem cell identity gene *WUSCHEL*, and down-regulation of CK signaling inhibitors (Fig. 3D, G), suggesting that plastid osmotic stress activates the CK/WUS feedback loop (Gaillochet et al., 2015; Ikeuchi et al., 2013) (top pathway, Fig. S6). The resulting imbalance in the cytokinin: auxin ratio may underlie callus formation in the *msl2 msl3* background, as supplementing seedlings with exogenous auxin robustly suppresses all mutant phenotypes (Fig. 3A, C) without affecting plastid osmotic stress (Fig 3B). In addition, the CK receptor AHK2 is required for efficient callus formation (Fig. 3E, F). *AHK2* is required to maintain WUS expression in the meristem in response to CK treatment in the CK/WUS feedback loop (Gordon et al., 2009). Cytokinin and auxin are involved in the production of all types of callus (Perianez-Rodriguez et al., 2014; Skoog and Miller, 1957), so alternative pathways to callus production cannot be ruled out.

Several previously identified callus-producing mutants in Arabidopsis and Helianthus also exhibit both up-regulated CK signaling and defects in meristem identity gene expression, including *tsd1* (Frank et al., 2002; Krupkova and Schmulling, 2009); EMB-2 (Chiappetta et al., 2006); and the *pasticcino* mutants (Faure et al., 1998; Harrar et al., 2003). These data suggest that, the CK-induced pathway, the wound-induced pathway and the meristematic identity pathway are interconnected pathways for callus production. Furthermore, the *msl2 msl3* double mutant is the only callus-producing mutant identified to date with a primary defect in plastid-localized proteins, adding a novel regulatory aspect to these known pathways, and suggesting that plastid osmotic stress is uniquely able to trigger these pathways. While we favor the model shown in Fig. S6, other interpretations are possible. Also consistent with these data is a model wherein the *msl2 msl3* lesions lead to down-regulation of *TSD1* and/or *TSD2*, or up-regulation of *WIND* genes, either of which could disrupt the CK/WUS loop.

### Plastid retrograde signaling is required for increased proliferation and the production of callus

Genetic lesions that reduce Pro biosynthesis significantly exacerbated callus production in the *msl2 msl3* background, an effect that can not be attributed to low levels of Pro itself (Figs. 1E, 4A, B, S4). Instead, the hyper-accumulation of ROS that results from blocking Pro biosynthesis is required (Fig. 5). Treatment with exogenous ROS scavenger TEMPOL as well as over-production of CSD2, a chloroplast-localized superoxide-scavenging enzyme, prevented or reduced callus formation. Plastid osmotic stress was required for ROS hyper-accumulation (Fig. 5A-C). In addition, the analysis of callus formation in higher order mutants established that ABA biosynthesis, the retrograde signaling protein GUN1, and the transcription factor ABI4 were required for callus formation and act downstream of ROS (Fig. 6, bottom pathway in Fig. S6). Because we were able to specifically suppress callus production, ROS accumulation, and impaired leaf development by increasing cytoplasmic osmolarity in the *msl2 msl3* background, this phenotype is very unlikely to be due to a loss of specific signaling by MSL2 or MSL3.

Multiple genetic links between plastid function and cell differentiation in the shoot have been previously described, and are often cited as evidence for retrograde signaling (Inaba and Ito-Inaba, 2010; Lepisto and Rintamaki, 2012; Lundquist et al., 2014). ROS accumulation has been documented in at least one other callus-producing mutant, *tsd2*/*quasimodo2* (Raggi et al., 2015). Whether ROS and/or an ABA/GUN1/ABI4 retrograde signaling pathway are employed to communicate plastid dysfunction to shoot development in any of these other mutants is not known. However, the application of pharmacological inhibitors of plastid development or function was used to demonstrate that leaf adaxial/abaxial patterning is regulated by plastid protein translation in a GUN1-dependent pathway (Tameshige et al., 2013), and the GUN1 pathway is required to facilitate the switch from leaf cell proliferation to expansion and differentiation (Andriankaja et al., 2012). Taken together with the results reported here, these data support previously proposed models wherein a variety of plastid dysfunctions are communicated to leaf development through similar or overlapping pathways that include GUN1 (Koussevitzky et al., 2007; Leon et al., 2012). ABI4 may function with GUN1 in this retrograde signaling pathway, or the reduction in *msl2 msl3* callus production in the *abi4-1* mutant background may result from indirect effects on sugar signaling or ABA biosynthesis.

We note that only a few of the genetic or pharmacological treatments that suppressed callus formation in the *msl2 msl3* mutant also suppressed the leaf developmental defects. Growth on exogenous auxin was the only treatment to completely suppress all of the mutant phenotypes, while preventing CK signaling, ABA biosynthesis, ROS accumulation, GUN1, or ABI4 function only partially rescued leaf defects (Figs. 3E-F, 6A-E). It is possible that these differences are due to genetic redundancy or to limited uptake or transport of TEMPOL. Alternatively, there may be two fundamentally different processes that respond to plastid osmotic stress in the *msl2 msl3* mutant: one that functions in the SAM during early development, and one that functions later on in the leaves. In support of the latter proposal, subjecting seedlings to mild hyperosmotic stress has been shown to prevent leaf cell proliferation (Skirycz et al., 2011).

### *msl2 msl3* mutants may provide a functional link between the CK/WUS feedback loop and plastid retrograde signaling in controlling cell differentiation at the plant apex

Our working model, illustrated in Fig. S6, is that the production of apical callus in *msl2 msl3* mutants operates through two non-redundant pathways: the CK/WUS feedback loop and a retrograde signaling pathway involving ROS, ABA, ABI4 and GUN1. While the data presented here establish that both of these pathways are required for callus formation in the *msl2 msl3* background, whether they operate completely independently or are interconnected is not yet clear. Here we present them as separate pathways; support for this comes from the fact that CK levels remain elevated in the *aba2 msl2 msl3* triple mutant (Fig. 3D), indicating that ABA biosynthesis is not upstream of the CK/WUS feedback loop. Furthermore, mutants that solely overproduce plastid ROS (Myouga et al., 2008) up-regulate CK signaling (To et al., 2004), or hyper-accumulate Pro (Mattioli et al., 2008) have not been reported to produce apical callus, implying that a combination of these signals is required.

We speculate that two-way communication between plastids and cell differentiation is essential to coordinate the developmental and functional state of the plastid with that of the cell within which it resides, and that it therefore necessarily involves multiple pathways. These results add yet another layer of complexity to the many regulatory pathways and feedback loops that govern dynamic cell identity decision-making at the plant shoot apex, and provide a foundation for future investigation into the relationship between meristem identity and plastid osmotic homeostasis in the SAM.

## MATERIALS AND METHODS

### *Arabidopsis thaliana* Mutants

The *aba2-1* (CS156), *abi4-1* (CS8104), *p5cs1-4* (SALK_063517), *p5cs1-1* (SALK_058000), *ahk2-2* (SALK_052531), *gun1-9*, and *msl2-3* alleles are in the Columbia*-0* background. The *msl3-1* allele is in the Wassilewskija background (Haswell and Meyerowitz, 2006). Derived cleaved-amplified polymorphic sequence genotyping (Neff et al., 1998) of the *gun1-9* allele was performed using oligos CGAACGACGAAAGATTGTGAGGAGGGTCT and CCTGCAAGCATTCAGAATCGCTGAAAAAGG, digesting with PstI. The *abi4-1* allele was genotyped with oligos TCAATCCGATTCCACCACCGAC and CCACTTCCTCCTTGTTCCTGC, digesting with NlaI. PCR genotyping of *msl*, *p5cs1*, and *aba2-1* alleles was performed as previously described (Sharma and Verslues, 2010; Szekely et al., 2008; Wilson et al., 2011).

### Plant Growth

Plants were grown on full-strength Murashige and Skoog (MS) medium (pH 5.7; Caisson Labs) with 0.8% (w/v) agar (Caisson Labs). NaCl, L-Pro (Sigma) and TEMPOL (Sigma) were added before autoclaving. For transfer assays, seed was surface-sterilized, sown on nylon mesh strips overlaid on 1X MS or 1X MS + 82 mM NaCl, and stratified for 2 d at 4°C before growth and transfer as described. All plants were grown at 23°C under a 16 h light regime, 130-160 μmol m^-1^ s^-1^.

### Microscopy

Confocal microscopy of ds-RED labeled non-green plastids was performed as in (Wilson et al., 2014). Bright-field images were captured with an Olympus DP71 microscope digital camera and processed with DP-BSW software. Apical meristems were ultra-rapidly frozen in a Baltec high- pressure freezer (Bal-Tec HPM010), excluding air with packing buffer (75 mM PIPES, pH 6.8, 50 mM sucrose). Samples were freeze substituted in 2% osmium/ acetone at −85°C for five days, slowly thawed to 25°C over 16 hours, and embedded in Spurrs resin. One μm sections for phase microscopy were stained 30 seconds with 1% Toluidine Blue O in 1% boric acid. Transmission electron microscopy was done on thin sections of tissue fixed for 2 h in 2% glutaraldehyde, post-fixed 90 minutes in 2% osmium tetroxide and embedded in Spurr’s resin. Sections were stained in uranyl and lead salts.

### ROS Detection and Quantification

Detection of H_2_O_2_ using 3,3’-Diaminobenzidine (DAB, Sigma) was performed as described (Wu et al., 2012) with the following modifications. Whole seedlings were incubated for 2 hrs in 1 mg/ml DAB prior to vacuum infiltration, incubated in the dark for an additional 12 hrs, then cleared with an ethanol series. An Amplex Red Hydrogen Peroxide/Peroxidase Assay Kit (Invitrogen) was used to measure H_2_O_2_ production in seedlings. For in vitro localization of O_2_^−^ with Nitrotetrazolium Blue chloride (NBT; Sigma), whole seedlings were vacuum-infiltrated with 0.1% (w/v) NBT in a 10 mM potassium phosphate buffer (pH 7.8) containing 10 mM NaN_3_. After 1-hour incubation in the dark at RT seedlings were cleared with an ethanol series. Quantification of formazan levels was performed as described in (Myouga et al., 2008).

### Subcloning and Transgenic lines

pENTR-MSL2 (Haswell and Meyerowitz, 2006) was used in a Gateway technology (Life Technologies) recombination reaction with pSCR:GW (Michniewicz et al., 2015) to create *pSCR:MSL2*.

### Quantitative Reverse Transcription-PCR

Quantitative reverse transcription-PCR was performed as previously described (Wilson et al., 2011). Primers used for gene expression analysis of *ACTIN* and *ARR* genes were previously described (Wilson et al., 2014; Zhao et al., 2010). The following primer pairs were used to amplify *WUS*(GCGATGCTTATCTGGAACAT and CTTCCAGATGGCACCACTAC) and *STM* (CAAATGGCCTTACCCTTCG and GCCGTTTCCTCTGGTTTATG).

## ACKNOWLEDGEMENTS

We thank Kelsey Kropp, Jeffrey Berry and the Proteomics and Mass Spectrometry Facility of the Donald Danforth Plant Science Center for technical assistance, Lucia Strader for *abi4-1* seeds and pSCR:GW, Ramanjulu Sunkar for pBIB-CSD2 and pBIB-mCSD2;,and Joanne Chory for *gun1-9* seeds. This research was funded by NSF MCB-1253103 and NASA NNX13AM55G.

## AUTHOR CONTRIBUTIONS

Study conception and manuscript preparation, MEW, ESH; conducting research and formal analysis, MEW, RHB, MM; funding acquisition, ESH.

## SUPPLEMENTARY FIGURE LEGENDS

**Fig. S1.**
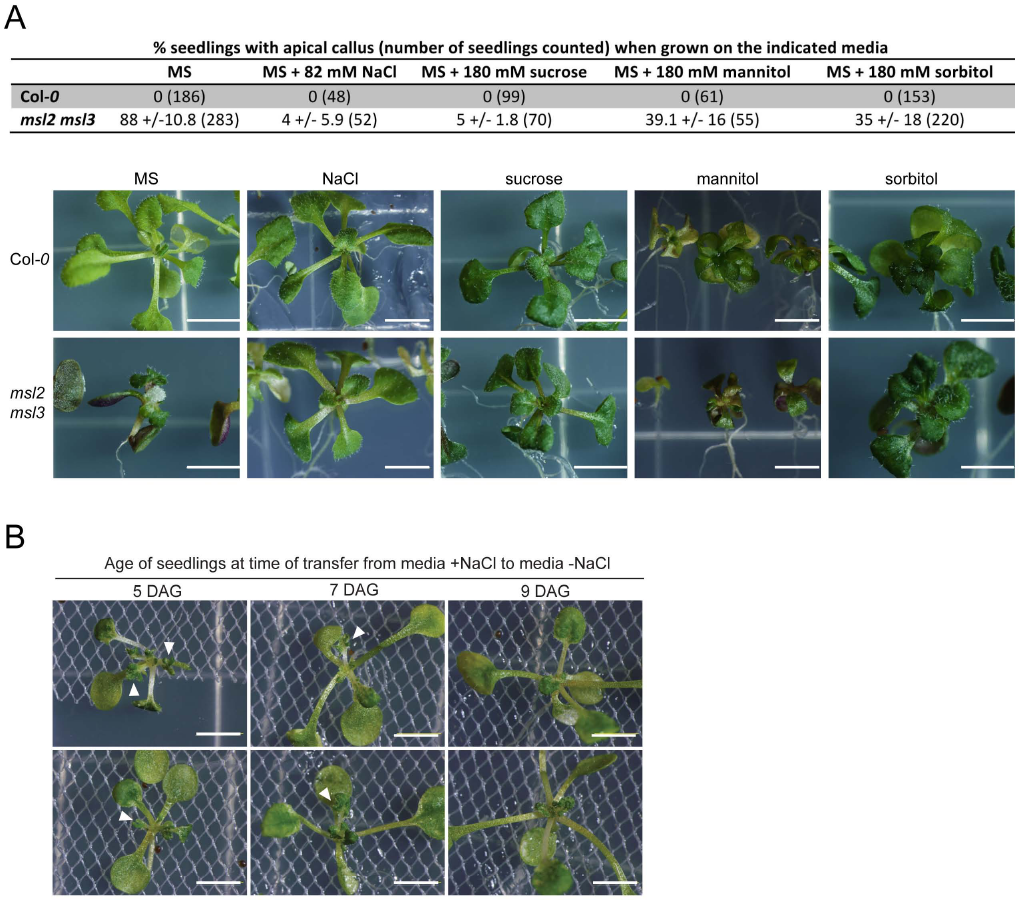
Additional analysis of callus production in *msl2 mls3* mutant seedlings. (A) Callus production is suppressed in *msl2 msl3* seedlings grown on MS supplemented with various osmotica. Top, percent of seedlings with visible apical callus after 3 weeks of growth on the indicated media. Bottom, individual seedling images. Scale bars are 5 mm. (B) *msl2 msl3* seedlings develop abnormal leaves after transfer from media containing osmotica to media without added osmotica. Bright field images of 21-day-old seedlings transferred at the indicated time points. Leaf notching is indicated by the white arrowheads.

**Fig. S2.**
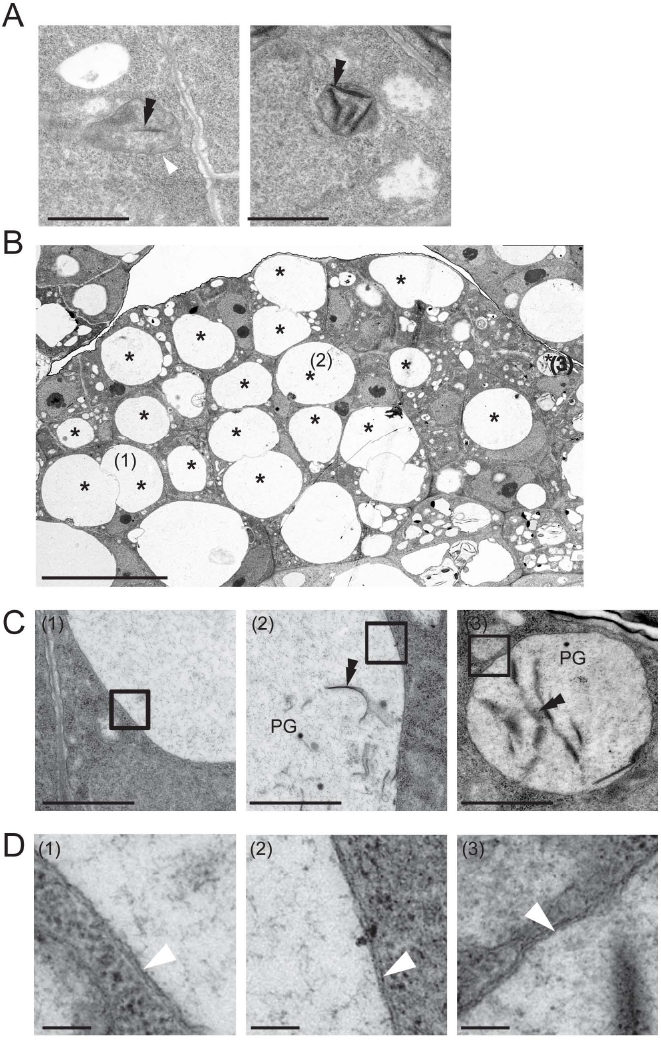
Additional transmission electron microscope (TEM) images of developing proplastids in the shoot apical meristem of 4-day-old wild type (A) and *msl2 msl3* mutants (B-D) seedlings. Size bar = 1 μm (A and C). (B) Proplastids are indicated (*), size bar = 10 μm. (C) Higher magnification images of the three proplastids marked with numbers in (B). Boxes in (C) are further magnified in (D) to show double membranes (white arrowhead). Size bar = 100 nm. Plastoglobules (PG), thylakoids (double black arrows).

**Fig. S3.**
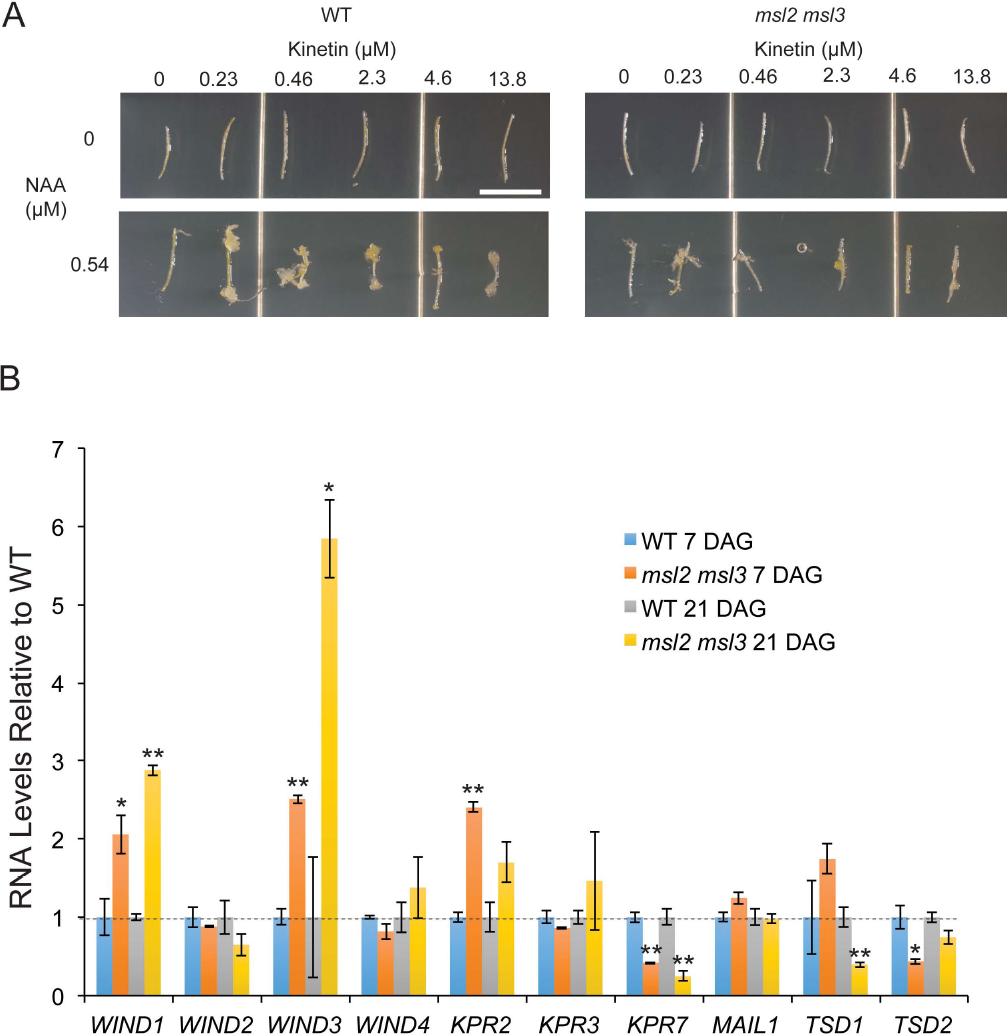
Characterization of callus production in *msl2 msl3* mutants. (A) Etiolated hypocotyl explants cultured on MS media containing the indicated concentrations of auxin (NAA) and cytokinin (kinetin). (B) Quantitative RT-PCR analysis of transcript levels of genes involved in callus production pathways in *msl2 msl3* mutants compared to the wild type. The average of three biological replicates (two technical replicates; n ≥ 25 seedlings each) is presented. Error bars indicate SEM, **P<0.01, *P<0.05 (Student’s *t*-test).

**Fig. S4.**
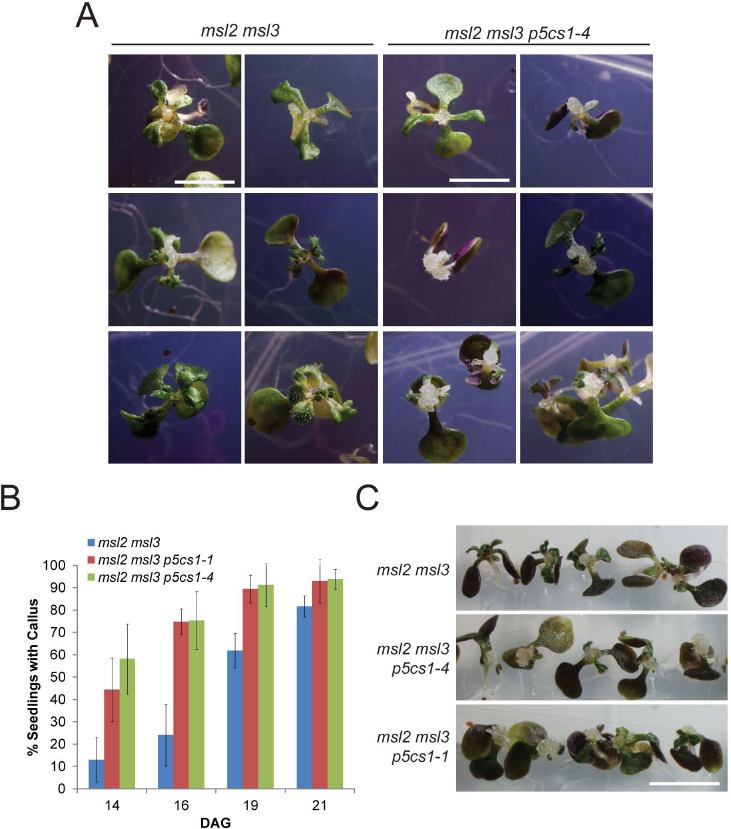
Two *p5cs1* alleles exacerbate callus production in the *msl2 msl3* background. (A) Bright-field images of 21-day-old *msl2 msl3* and *msl2 msl3 p5cs1-4* mutants. Size bars, 5 mm. (B) Callus production at various time points after germination in *msl2 msl3*, *msl2 msl3 p5cs1-4*, and *msl2 msl3 p5cs1-1* seedlings. 5 plates of seedlings, n > 10 seedlings per plate, were analyzed per time point. (C) Bright-field images of *msl2 msl3 p5cs1-4* and *msl2 msl3 p5cs1-1* triple mutant seedlings at 21 DAG. Size bars, 5 mm.

**Fig. S5.**
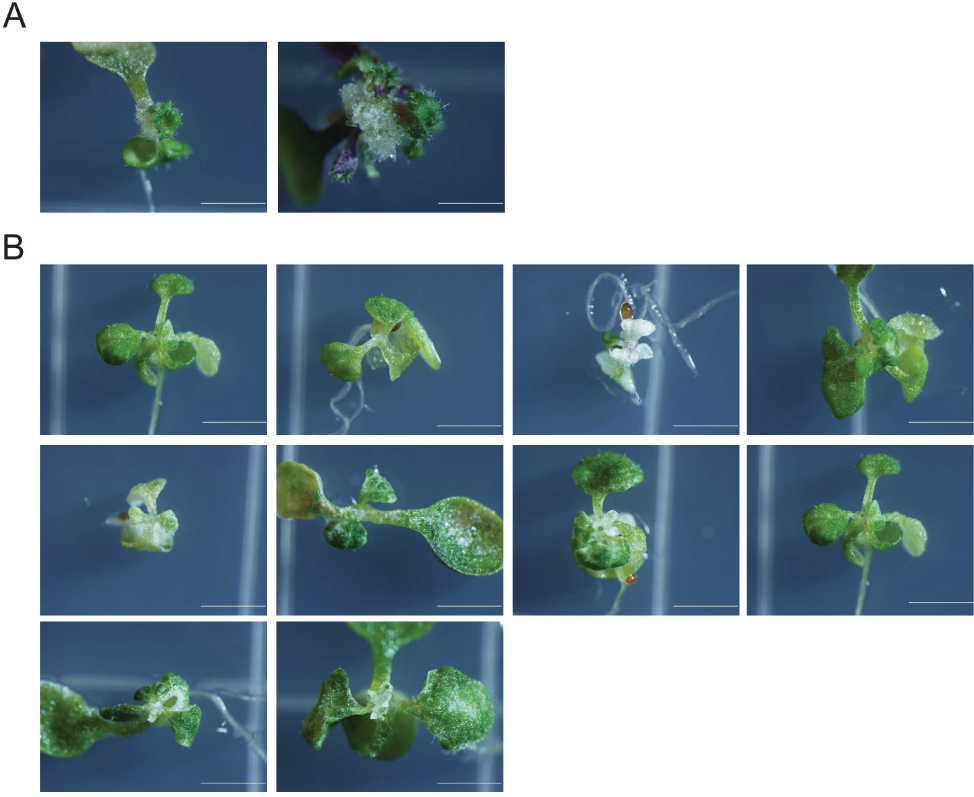
Additional images of *msl2 msl3 gun1-9 +/−* (A) and *msl2 msl3 gun1-9* triple mutant (B) siblings. Seedlings were grown for 21 days on solid media before imaging. Size bars, 2.5 mm.

**Fig. S6.**
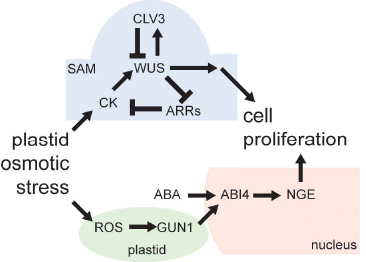
Working model. These results are consistent with a working model wherein plastid osmotic stress impacts both a feedback loop regulating cell proliferation in the SAM (based on (Gaillochet et al., 2015) and (Ikeuchi et al., 2013) and a plastid ROS-dependent retrograde signaling pathway (Leon et al., 2012). Plastid osmotic stress leads to increased proliferation at the SAM and the production of callus through increased CK levels and *WUS* expression and through an unknown nuclear transcription program triggered by GUN1, ABA, and ABI4.

## SUPPLEMENTARY METHODS

### *In Vitro* Callus Production

Induction of *in vitro* callus production was performed as previously described (Iwase et al., 2011). Etiolated hypocotyls were excised from mutant and wild type seedlings 4 days after germination. Mutant and wild type explants were cultured for 30 days side-by-side on MS medium supplemented with Gamborg’s vitamins, 2% glucose, 0.05% 2-(N-morpholino)ethanesulfonic acid (MES), pH 5.8 with KOH, 0.8% phytoagar, and the indicated concentrations of NAA and kinetin.

### Quantitative RT-PCR

Quantitative reverse transcription-PCR was performed as previously described (Wilson et al., 2011). Primers used for gene expression analysis of *KPR and WIND* genes were previously described (Iwase et al., 2011; Schiessl et al., 2014). The following primer pairs were used to amplify *TDS1* (AGGCACTGCGGAATCTAGCAA and CTGGTCTGATTGTGGAAGGTCG) and *TDS2* (CAGGTCAGCATCTCCTATGTCT and CCAAGATAGTTCTCACACCAGC) and *MAIL1* (CTTGATCCTAGCAAGATCACTTGG and TGCCTCAGACACCTATCTGGA).

